# Emergence of a contrast-invariant representation of naturalistic texture in macaque visual cortex

**DOI:** 10.1101/2025.01.03.631258

**Authors:** Gerick M. Lee, Najib J. Majaj, C. L. Rodríguez Deliz, Lynne Kiorpes, J. Anthony Movshon

## Abstract

Sensory stimuli vary across a variety of dimensions, like contrast, orientation, or texture. The brain must rely on population representations to distinguish changes in one dimension from changes in another. To understand how the visual system might extract separable stimulus representations, we recorded multiunit neuronal responses to texture images varying along two dimensions: contrast, a property represented as early as the retina, and naturalistic statistical structure, a property that modulates neuronal responses in V2 and V4, but not in V1. We measured how sites in these 3 cortical areas responded to variation in both dimensions. Contrast modulated responses in all areas. In V2 and V4, the presence of naturalistic structure both modulated responses and increased contrast sensitivity. Tuning for naturalistic structure was both strongest and most dispersed in V4. We measured how well populations in each area could support the linear readout of both dimensions. Populations in V2 and V4 could support the linear readout of naturalistic structure, but in V4, this readout was more robust to variations in contrast.

**Significance Statement:** To support flexible behavior, the brain must simultaneously represent different stimulus dimensions. Single neurons are typically modulated by multiple dimensions, and so cannot distinguish them - they must be extracted by decoding neural populations. We studied neuronal responses in three cortical visual areas - V1, V2, V4 - using texture images varying in both contrast and naturalistic image structure. We used population decoders to read out each dimension. In all areas, contrast was well decoded independently of image structure. On the other hand, V1 could not decode image structure independent of contrast, while V2 and V4 could. V4 decoding was greatly superior, because the selectivity of individual sites for texture was more diverse than in V1 or V2.

---

The responses of sensory neurons are modulated by multiple stimulus dimensions, so a change in neuronal response does not unambiguously represent a change in any particular dimension. To support the readout of multiple stimulus dimensions, there must be neurons that are diverse in their stimulus selectivity – a population of identically tuned neurons would fare no better than one.

Consider a case in point. The primates reading this paper can do so across a variety of contrast and lighting conditions, from backlit digital displays with strongly contrasting black text set against a bright white background, to poorly printed paper copies in a dim room. This “contrast-invariant” capacity requires the perceived identity of a word to remain stable across different lighting conditions. It also exemplifies the larger computational problem – the brain must represent a large number of sensory dimensions (e.g., contrast, stimulus identity), and must flexibly support the readout of any of these dimensions, based on the perceptual demands of the moment (1, 2).

Here, we focused on two stimulus dimensions. One, visual contrast, is explicitly encoded by neurons in the retina (3, 4). The second concerns the structure of natural images. A characteristic feature of these images is the presence of structures that reflect spatial correlations, like edges, lines, and contours (5). Texture images that contain this “naturalistic” structure often drive neural activity in visual areas V2 and V4, more strongly than “noise” textures, matched in orientation and spatial frequency content (6–8). The presence of naturalistic structure does not modulate neuronal activity in V1 (6, 8–10).

We recorded the responses of neural populations in areas V1, V2, and V4 of awake macaque monkeys during passive fixation. We presented texture images, with and without naturalistic structure, at a range of contrasts. We measured how contrast and texture were encoded at individual multiunit sites. Sites in all areas were modulated by contrast. Sites in V2 were often also modulated by naturalistic structure; sites in V4 more so. In V4, this modulation was higher on average, and was more dispersed. In V2 and especially in V4, encoding of contrast and texture intersected: contrast sensitivities were higher in response to naturalistic textures than to noise textures. We measured how well populations in all 3 areas could separately encode both contrast and naturalistic structure. As expected, the V1 population encoded only contrast, and could not distinguish naturalistic structure. The V2 population encoded both variables, but the representations of naturalistic structure and contrast in V2 were weaker than in V4. In V4, we found a more robustly contrast-invariant representation of naturalistic structure which could be extracted using a simple, biologically-plausible mechanism, and which depended on sites being heterogeneous in their tuning for naturalistic structure.

## Results

Neuronal responses in V2 and V4 are often modulated by the presence of naturalistic structure within images, while responses in V1 are not. To understand how this modulation is affected by changes in contrast, we used data from 58 multiunit sites from V1, 120 sites from V2, and 108 from V4. We presented naturalistic textures, and their spectrally matched noise counterparts, at different contrasts (Fig. 1).

**Fig. 1.**
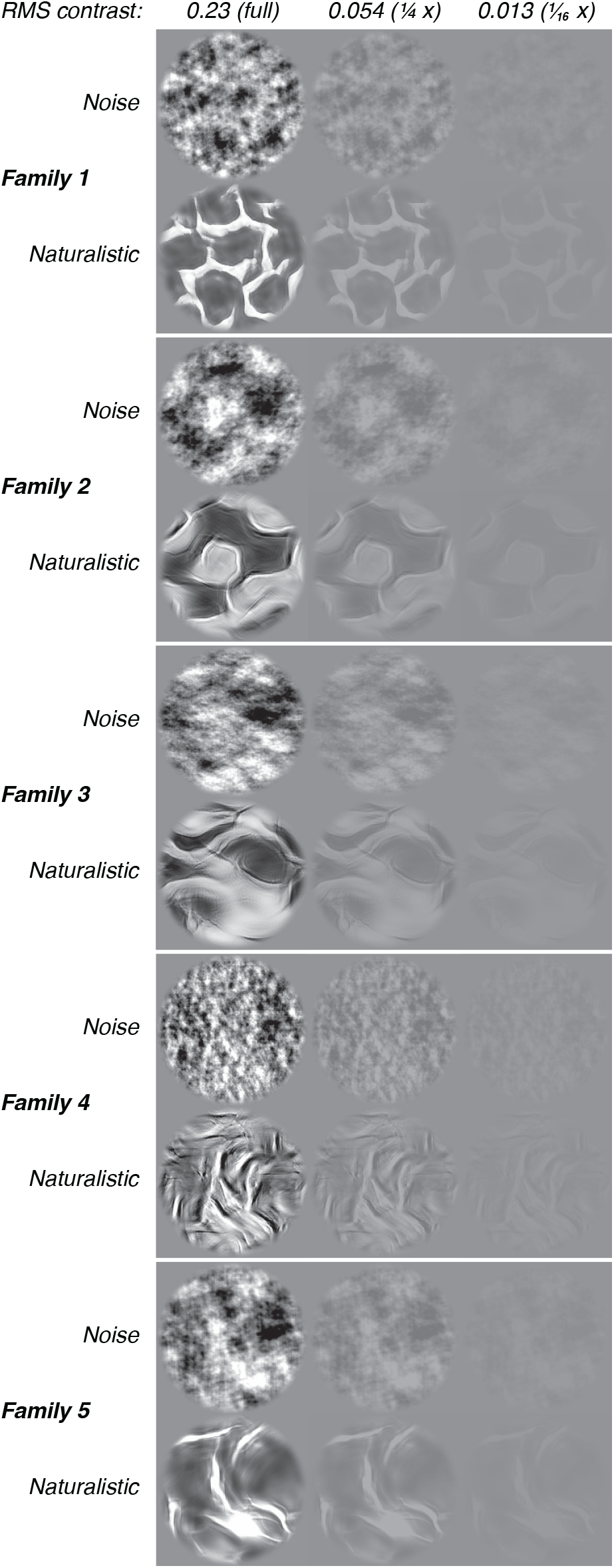
Synthetic texture images as used in our experiments. We synthesized “noise” (above) and “naturalistic” (below) textures to match the average spectral statistics of natural images. We used 5 natural images to generate the different texture “families” shown here. We displayed these images at 7 total contrasts, 3 of which are shown here (values at the top are the average RMS contrast across all textures at that contrast). Note the presence of sharp, continuous elements (e.g., edges, lines, contours) in the naturalistic textures; these elements are largely absent from the noise textures. The difference between naturalistic and noise textures is perceptible at all 3 contrasts, even near the threshold for seeing.

### Characterizing contrast sensitivity

In all 3 areas, contrast affected both the magnitude and onset latency of responses (Fig. 2A). Responses in V1 were not modulated on average by the presence of naturalistic structure. The average response in V2 and V4 to naturalistic textures was stronger than the response to spectrally-matched noise textures. In V1 and V2, the transient onset response to high contrast was stronger for noise than for naturalistic textures. In V2 and V4, low contrast stimuli evoked an onset response which was weaker and slower than high contrast stimuli. Yet even at low contrasts, naturalistic textures evoked a larger response than noise textures.

**Fig. 2.**
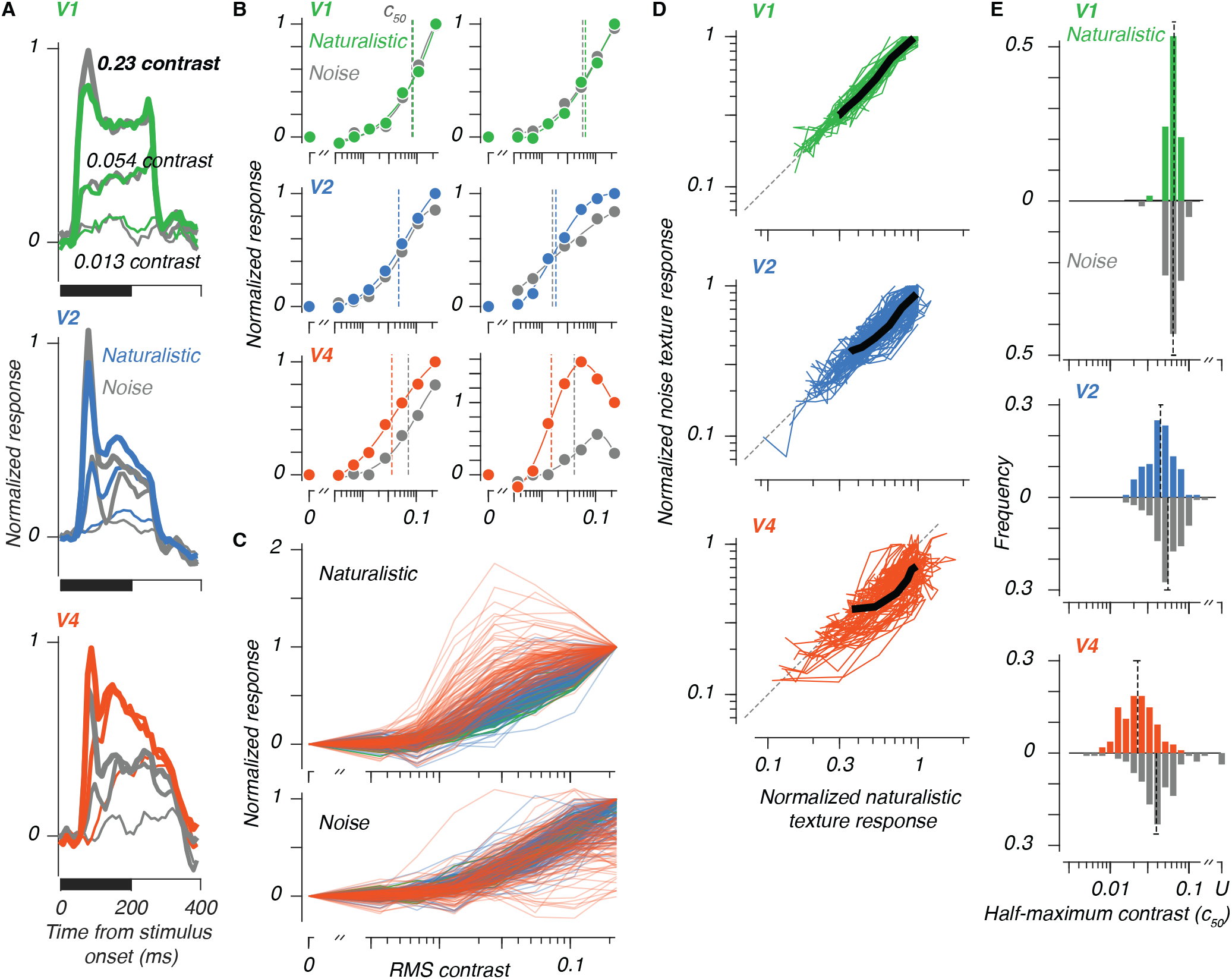
Multiunit neural responses to texture images at different contrasts. A: Post-stimulus time histograms, each normalized to the maximum response, then averaged across all responsive sites, to naturalistic and noise textures at 3 different contrasts (the same contrasts shown in Fig. 1). The mean latencies of the sites included in these mean traces at full contrast were 47, 56, and 74 ms. B: Responses for six example sites, for the first 200 ms of the visually evoked response. Responses are normalized to the naturalistic texture response at maximum contrast. Points show average responses to naturalistic (full color) and noise textures (gray), solid lines show model fits, using the model of Peirce (11). Half-maximum contrast values for naturalistic and noise textures (*c*_50_) are marked with vertical dashed lines (colored and gray, respectively). C: Contrast responses to texture stimuli for all sites. The upper panel shows responses to naturalistic textures, the lower panel shows responses to noise textures. Both sets of curves are normalized to the largest response at maximum contrast for each site, whether it was evoked by naturalistic or noise stimuli. D: Contrast responses for all sites, normalized to the largest response at maximum contrast (as in C). Black lines show geometric means. E: Distributions of half maximum contrast values (*c*_50_) for each area and stimulus type. Vertical dashed lines and numerical values show medians. The horizontal line at the extreme end of each dashed line shows the 95% confidence interval around the median. For some sites in V4, a half maximum contrast could not be fit for the data to noise textures, because the response was too weak or variable. These “undefined” sites (U) are offset to the right, and were not used to compute medians. We used the *c*_50_ values as a proxy for contrast threshold.

We measured the relationship between contrast and response at the level of individual sites (Fig. 2B). In V1, responses increased monotonically with contrast. Responses to naturalistic and noise textures were comparable at all contrasts. Responses in V2 also increased monotonically with contrast. At higher contrasts, both example sites responded more to naturalistic textures than to noise textures. In V4, we saw two changes relative to V1 and V2 – first, we noticed a divergence between sites’ contrast sensitivity measured using naturalistic textures, versus their noise texture evoked contrast sensitivity. As a result, while V4 response magnitudes varied with contrast, a selectivity for naturalistic texture was visible across a wide range of contrasts. Second, we noticed an increased prevalence of supersaturating contrast-response relationships, typified by the bottom right example in Fig. 2B.

To quantify the contrast response function across all sites in our sample, we fit responses from all sites with the model of Peirce (11), which uses a modified Naka-Rushton function to relate response and contrast. The model includes a parameter which allows it to capture these supersaturating contrast-response relationships. We used the model fits to measure the contrast at which responses reached half of their maximum (*c*_50_), which we take as a proxy for contrast threshold, the inverse of which is contrast sensitivity. Lower *c*_50_values indicate higher contrast sensitivities. In the examples shown in Fig. 2B, contrast sensitivity was higher in V2 and V4 than in V1.

Figure 2C shows the contrast response functions for all sites, in response to naturalistic (above) and noise textures (below). For naturalistic textures, contrast response functions taken in V4 (orange traces) were typically to the left of contrast response functions taken in V1 and V2 (green and blue traces, respectively), suggesting that V4 sites had higher contrast sensitivity for naturalistic textures. This difference was less evident for noise textures. Supersaturating contrast response functions were also more prevalent in V4 than in V1 or V2, especially for responses to naturalistic textures.

Figure 2D plots the normalized response for each site to noise against the response to naturalistic texture. The traces for V1 and V2 were typically near or along the diagonal, meaning that responses to the two texture types were roughly equal at all contrasts. In V4, the curves lie below the diagonal, meaning that the responses to naturalistic textures were typically stronger than to noise, even at intermediate contrasts.

We compared half maximum response (*c*_50_) values (taken from Peirce’s model), measured separately for naturalistic and noise textures, for all areas (Fig. 2E). Taking these *c*_50_values as measures of contrast threshold, the figure reveals two main trends in the data. First, contrast thresholds fell (i.e., sensitivity increases) from V1 to V2, and again from V2 to V4. Second, the difference in contrast threshold between responses to naturalistic and noise textures was largest in V4, smaller in V2, and absent in V1. For V2 and V4, thresholds were lower for naturalistic textures than for noise.

We compared interareal contrast thresholds via permutation ANOVA, followed by pairwise permutation tests (permutation ANOVA (12), naturalistic: *F* = 56, *p <* 0.001; noise: *F* = 11, *p <* 0.001, pairwise permutation tests: V1-V2 noise texture difference: *p <* 0.005; all other comparisons *p <* 0.001). We tested the differences between naturalistic and noise texture evoked contrast thresholds via pairwise permutation tests (V1: *p >* 0.05, V2: *p <* 0.001, V4: *p <* 0.001). In words: other than the naturalistic-noise difference in V1, these effects were statistically significant.

We also measured the fraction of supersaturating contrast response functions, using the fits to the model of (11). As shown above (Fig. 2B and C), all V1 responses to naturalistic and noise textures increased monotonically with contrast. In V2, 4 of 120 sites (3%) had a supersaturating contrast response – 3 to both texture types, 1 to noise textures. In V4, 51 of 108 sites (47%) had a supersaturating contrast response – 11 to both texture types, 14 to naturalistic textures, and 26 to noise textures. These supersaturating sites typically had lower *c*_50_values than monotonic sites: the geometric mean naturalistic texture evoked *c*_50_in V4 was 0.024 for monotonic sites, versus 0.018 for supersaturating sites. In V2, these values were 0.044 and 0.021, respectively.

### Single site encoding of naturalistic structure and contrast

Having characterized the relationship between stimulus contrast and neural response, we wanted to measure the selectivity of each site for naturalistic image structure, first between full contrast naturalistic and noise texture pairs, then for all other contrast pairs.

We used the separation between naturalistic and noise texture response distributions (Fig. 3A) to measure texture selectivity. Measured at full contrast, this comported with prior observations – on average, sites in V1 barely preferred noise textures over naturalistic ones (mean d’ [95% CI]: -0.07 [- 0.11, -0.04]); sites in V2 and V4 preferred naturalistic textures to noise (V2 mean d’ [95% CI]: 0.15 [0.10, 0.19]; V4: 0.50 [0.41, 0.59]). We compared each of the 3 distributions, and found a significant effect of area on selectivity (permutation ANOVA: *F* = 22, *p <* 0.001). Followup comparisons confirmed that V2 was significantly more selective than V1 (mean difference: 0.21; permutation test, *p <* 0.001). V4 was significantly more selective than V2 (mean difference: 0.36, permutation test, *p <* 0.001), and V1 (mean difference: 0.57, permutation test, *p <* 0.001). These relationships largely agree with prior observations, with the exception that the V4-V2 difference reported here is larger than in our previous study, based on data from the same series of recordings but not from the same sessions (8). In those sessions, we recorded neuronal responses and reported d’ values for 35 texture families, at maximum contrast, and at a fine spatial scale. In these sessions, we used only 5, at several contrasts, at a coarser spatial scale, chosen on the basis of their ability to elicit a strong behavioral response to the difference between naturalistic and noise textures (see Fig. 1A & B in Lee et al., 2024 (8)). These stimulus differences likely account for the differences in d’, by decreasing selectivity in V2 and enhancing selectivity in V4. We suspect that the 2.7-fold lower spatial frequencies in the stimuli used in this study are the cause, but we cannot be sure without further experiments. More generally, the quantitative differences across measurement conditions will inevitably depend on the particulars of the stimulus set; the qualitative effects that are the focus of this analysis do not depend on these particulars.

**Fig. 3.**
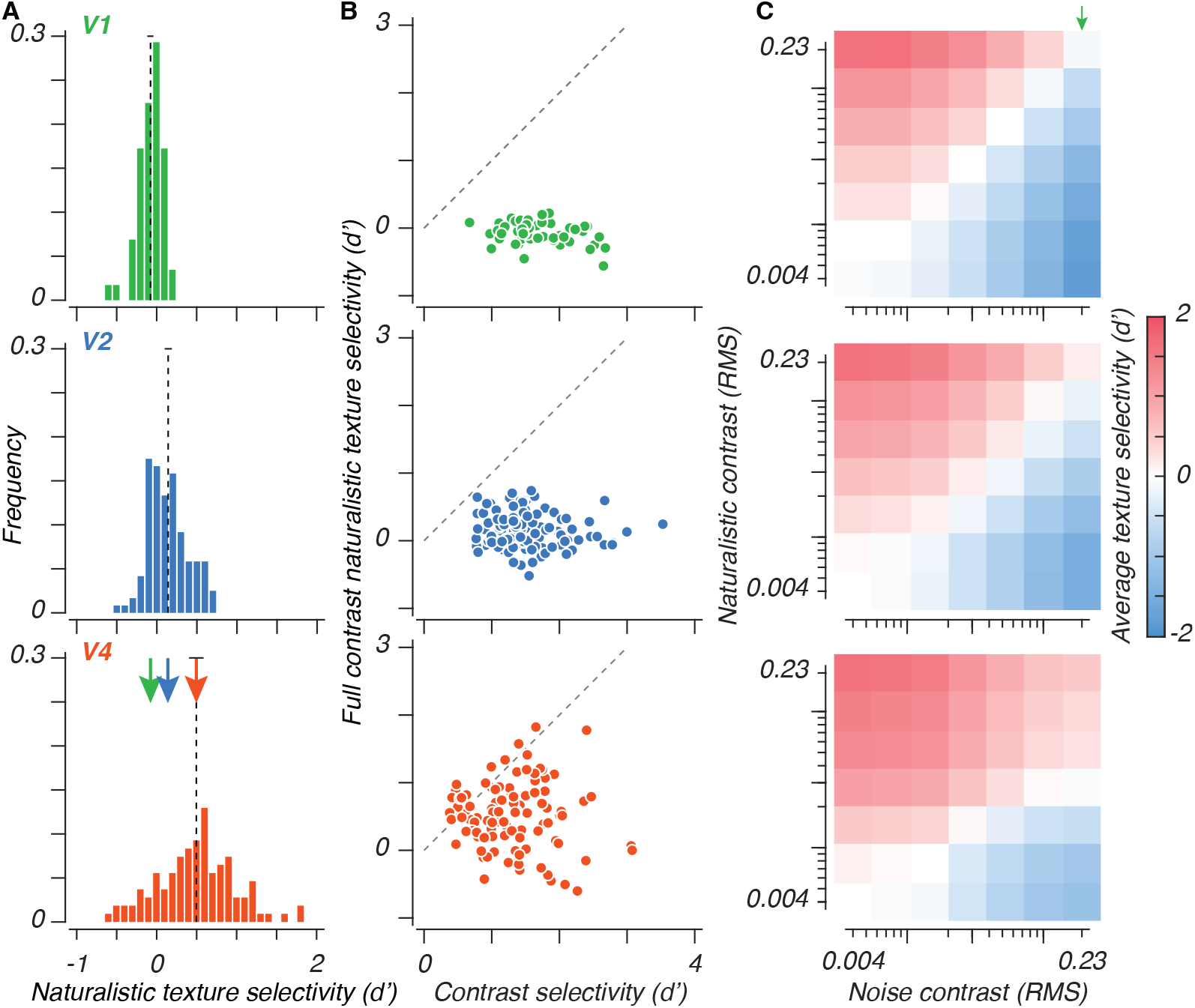
Single site texture selectivity. A: Distributions of single site texture selectivity, measured between the responses to full contrast naturalistic and noise texture images. Dashed lines and arrows show means. For reference, the selectivities for the example sites in Figure 2B were: 0.02 and 0.04 (V1), 0.32 and 0.22 (V2), 0.39 and 0.62 (V4), respectively. B: Contrast selectivity, measured as the d’ separating all responses to full contrast, from all responses to the lowest (non-zero) contrast, versus full contrast texture selectivity, as measured in panel A. Note that the identity lines (dashed gray) are vertically offset from the center of the plot. C: Mean single site texture selectivities, measured between responses to naturalistic and noise textures for all possible contrast pairs. The full-contrast selectivities shown in panel A are at the top right corner, indicated with a green arrow. In V1 and V2, selectivity is dominated by contrast – most heatmap cells above the identity diagonal are red, while most below are blue. In V4, the interaction between contrast and image structure is shown by the larger red region, showing contrast pairs for which naturalistic textures evoke stronger responses.

Having confirmed that V2 and V4 were both sensitive to the presence of naturalistic structure within texture images, we related each site’s selectivity for naturalistic texture to its corresponding contrast selectivity – the difference in response between full contrast stimuli (including both naturalistic and noise textures), and the lowest (non-zero) contrast (Fig. 3B). In V1, we observed a broad distribution of selectivity for contrast, while texture selectivity was negligible. Sites in V2 and V4 varied in their tuning for both contrast and texture. There was a large degree of variation along both axes in V4 – contrast and texture selectivity were heterogeneously distributed.

We computed the mean naturalistic-noise texture selectivity for all possible contrast pairs. Each heatmap in Figure 3C shows the texture selectivity for each of the 49 possible contrast pairs (7 naturalistic contrasts by 7 noise contrasts). The top right corner of each heatmap shows the same full contrast comparison as Figure 3A. Values along the identity diagonal – plotted from bottom left to top right – show the 7 matched contrast comparisons. Off-diagonal values show selectivities which reflect an area’s combined modulation by contrast and texture.

Responses in V1 were not selective for naturalistic structure – values were close to zero along the identity diagonal. Responses in V1 were selective for contrast – values for textures differing in contrast (off-diagonal) were large and reflected the ratio of contrasts (i.e., the difference in log contrasts) – selectivities were largest for pairs with the largest contrast ratio. Selectivities were negative for all contrast pairs below the identity diagonal: sites responded more vigorously to higher contrast noise textures than to lower contrast naturalistic textures.

As in V1, responses in V2 were also driven by contrast: for all off-diagonal contrast pairs, selectivity reflected the difference in contrast, not texture. As in V1, this included negative selectivities for all pairs below the identity diagonal. But in V2, responses along the identity diagonal were positive (red). While V2 is selective for naturalistic texture, our results suggest that contrast, not texture, is the predominant determinant of activity in V2.

As in V2, responses in V4 were selective for naturalistic structure. But in V4, naturalistic structure was more important than contrast for a number of unequal contrast pairs – demonstrated by the presence of a number of positive selectivities (red cells) below the identity diagonal.

Even in V4, selectivities were negative if the contrast ratio between lower contrast naturalistic and higher contrast noise was great enough (e.g., the bottom right corner of Figure 3C). Taken at face value, this result would make a paradoxical prediction about texture perception, best demonstrated by Figure 1 – specifically, the rightmost (lowest contrast) naturalistic textures, and leftmost (highest contrast) noise textures. For these comparisons, a typical human observer can readily identify which is the more naturalistic. If behavioral texture discrimination resulted from the simple average of single site activity, as seen in Figure 3C, observers would not only be unable to correctly distinguish low contrast naturalistic textures from full contrast noise (i.e., chance performance), but also they would consistently perform *below* chance, and mislabel high contrast noise textures as naturalistic. A glance at Figure 1 demonstrates that this is not consistent with human experience. We therefore wanted to consider how ensembles of neurons might be able to disambiguate contrast from naturalistic structure.

### Extracting a contrast-invariant population code for naturalistic texture

We trained two linear classifiers, one to discriminate contrasts (using both naturalistic and noise textures), and another to discriminate naturalistic from noise textures (using responses to all pairs of contrasts and all texture families). We trained and tested each classifier on multiple randomly sampled populations of 32 sites. At the population level, contrast discrimination was comparable between V1, V2, and V4 – all off-diagonal regions of the heatmaps are colored to indicate that contrasts are correctly discriminated by the populations regardless of naturalistic structure (Fig. 4A).

**Fig. 4.**
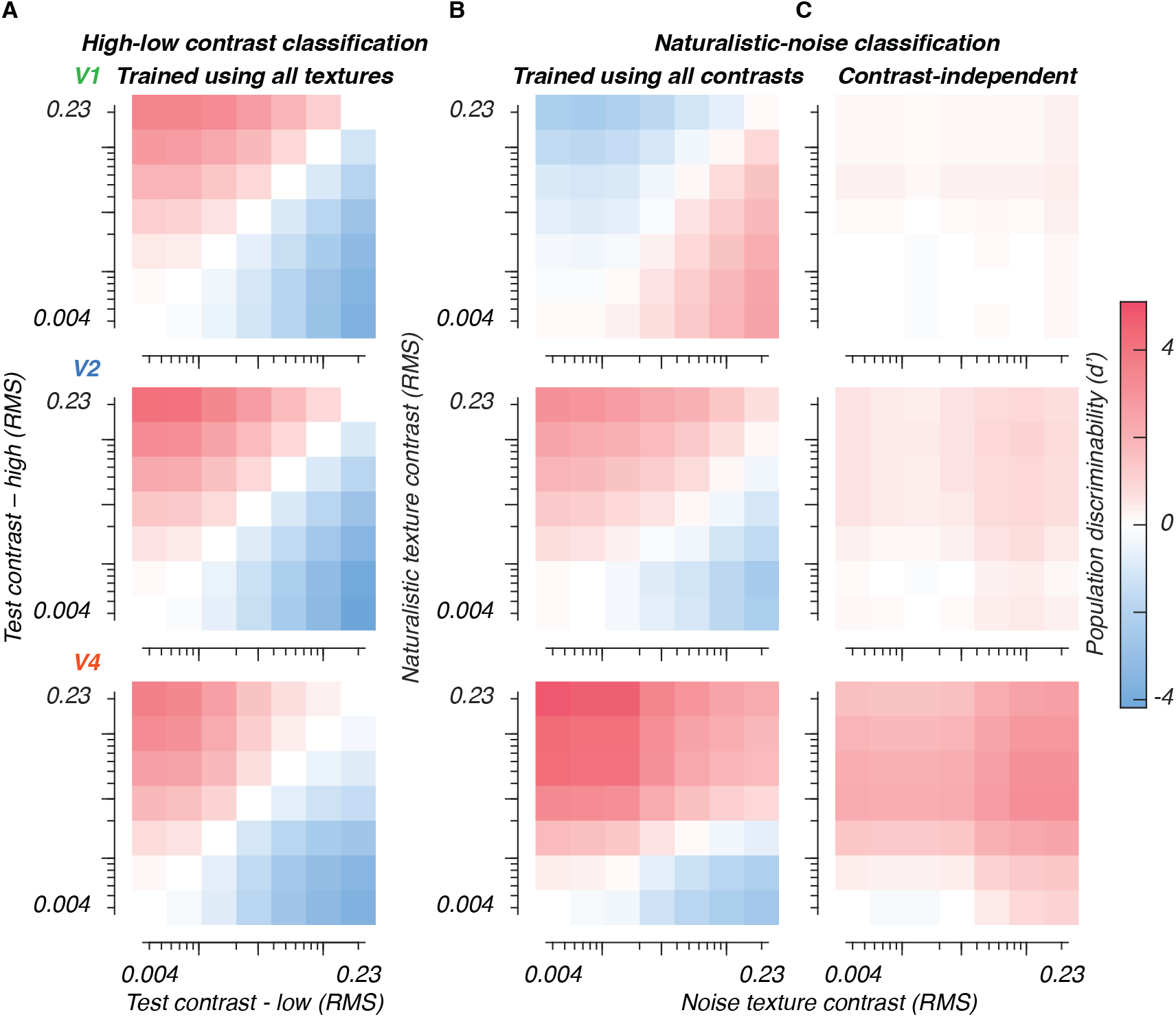
Population contrast and texture discriminabilities. A: Mean contrast discriminability, based on populations of 32 sites. We trained a classifier to discriminate high from low contrast textures, using neural responses to both naturalistic and noise textures. We tested performance separately for each contrast pair (using held-out data). Perfect performance for this decoder would yield a heatmap that is red above the identity diagonal and blue below it – very much as shown. B: Mean naturalistic-noise texture discriminability, based on populations of 32 sites. We trained a classifier to discriminate naturalistic textures from noise textures, using neural responses to all contrasts and all families. We tested performance separately for each contrast pair. Perfect performance for this decoder would yield a heatmap that is everywhere red. Plainly performance is incorrect (blue) in many cases. C: Mean naturalistic-noise texture discriminability, based on populations of 32 sites. We used the Gram-Schmidt process to obtain a third classifier: a naturalistic-noise classifier, orthogonal to the contrast classifier in panel A. We used this contrast-independent classifier to test performance separately for each contrast pair. Perfect performance for this decoder would again yield a heatmap that is everywhere red. Performance is generally excellent for V4, and often correct though much weaker in V2 and V1.

We used the second classifier to measure discriminability for naturalistic versus noise textures across all contrast pairs (Fig. 4B) – a correct classifier for naturalistic texture should yield positive classification performance for all contrast pairs, i.e. all heatmap cells should be red. V1 could not reliably discriminate naturalistic textures from noise, and instead often mislabeled low contrast stimuli as naturalistic textures. This was a result of the noise-preferring onset transient visible in Figure 2A – excluding this transient from the analysis window reduced performance to near zero. This shows that the population did not reliably represent naturalistic statistics, leaving the classifier with only the nuisance dimension of contrast. Populations in both V2 and V4 supported texture discrimination for many contrast pairs. Yet as was the case for the single-site average (Fig. 3B), the representations of texture and contrast were conflated, leading to below-chance performance for contrast pairs that contained low contrast naturalistic textures and high contrast noise (blue zones at the bottom right corner of both heatmaps). A suitable measure of classifier performance in this case is the mean d’ for all values below the identity diagonal, i.e. for all cases in which the naturalistic texture was of lower contrast than the noise texture. This “contrast-invariant discriminability” was positive for V1 (1.2), and negative (i.e. incorrect) for V2 (-0.79) and V4 (-0.25).

If neurons in a population differ from one another in their tuning for each dimension, information can be read out by orthogonalizing the representation of the nuisance dimension (in this case, contrast). If all sites are identically tuned for a pair of stimulus dimensions, the system is underdetermined, and the dimensions cannot be differentiated. Our measurements in V2 and V4 suggested that tuning for contrast and texture was heterogeneous, especially in V4 (Fig. 3B). We therefore expected that a correctly chosen linear classifier could leverage the diversity in tuning seen in the V4 population to support contrast-invariant texture discrimination.

The two classifiers we trained – the contrast discriminant and the texture discriminant – were both vectors in the same population response space. As a result, we were able to use standard linear methods (Gram-Schmidt process) to project the contrast discriminant axis (Fig. 4A) out of the texture discriminant axis (Fig. 4B). This resulted in a third discriminant – a new texture discriminant axis *orthogonal* to the contrast discriminant. We used this contrast-independent axis to again measure how well each area could discriminate naturalistic textures from noise (Fig. 4C). Performance in V1 was close to zero for all contrast pairs, suggesting that the original learned representation of naturalistic structure was primarily driven by stimulus contrast, not by texture statistics. In V2, performance was positive for most contrast pairs, the exception being pairs in which both stimuli were of low contrast. In V4, performance was again positive for most pairs, except where both stimuli were low in contrast. Notably, all three areas show correct performance for almost all contrast pairs (all cells are to some degree red). Using the same figure of merit as above, we now observed robust contrast-invariant discriminability in V4 even for cases in which contrast and naturalistic statistics were in conflict (mean d’ of heatmap cells below the identity diagonal = 1.6). Performance was weakly positive in V2 (mean d’ = 0.45), and weaker still in V1 (mean d’ = 0.11). To determine whether contrast invariance in V4 was dependent on sites with a supersaturating contrast-response relationship, we computed classifier performance with all supersaturating sites excluded. Performance was highly similar – suggesting that contrast-invariant performance did not depend on the inclusion of sites with supersaturating responses.

Comparison of Figure 4B and C shows that the orthogonalized classifier performed worse for contrast pairs featuring high contrast naturalistic, and low contrast noise textures, but its performance was superior for matched contrasts, and for contrast pairs where the noise texture was higher in contrast than the naturalistic one. Critically, this shift in performance was possible using exclusively linear operations. Our results thus demonstrate that a biologically-plausible, contrast-invariant representation of naturalistic texture progressively emerges through the first areas of the ventral stream.

### Heterogeneous texture selectivity supports contrast invariance

The sites we recorded in V2 and V4 often encoded both contrast and texture. As a result, the separable population representations of these two dimensions must take the form of separate subspaces within the same population response space. That is, there must be orthogonal axes within this population response space which uniquely represent either contrast or texture, even if the individual neurons in the population are modulated by both. We supposed that the usefulness of these separate axes for decoding should depend on heterogeneity in the population.

To relate population contrast invariance with single-site tuning properties, we repeated the contrast-invariant classification shown above (Fig. 4C), but using populations of 8 sites. Smaller populations will contain fewer overlapping sites, and their tuning will be more variable. This allowed us to better measure the impact of heterogeneity on population contrast invariance. For each subsample, we measured contrast-invariant discriminability, and the mean and standard deviation of selectivities for texture and contrast. We then used a multivariate regression to determine which of these factors best predicted contrast invariance (13).

As a consequence of the smaller population size, overall classifier performance was reduced (Fig. 5A): contrast-invariant discriminability was 0.04 in V1, 0.29 in V2, and 0.77 in V4. We fit a four-factor model to measure the standardized partial regression coefficients *β*. These values represent the effect of each variable on contrast-invariant performance when each variable is normalized to a standard (Z-transformed) scale. The larger the *β*, the stronger the relationship between the predictor and contrast-invariant performance. As Table 1 shows, in all three areas, that the best predictor of contrast invariance was the standard deviation in texture selectivity. The relationship between contrast invariance and texture heterogeneity is shown in Figure 5B (standardized relationships for all four predictors are shown in Fig. S1). Contrast invariance is typically smallest in V1, and largest in V4, as already noted. But observe that both within and between areas, the greater the texture heterogeneity, the more robust the contrast invariance. Note also that subsamples taken from different areas which match in heterogeneity have similar values of contrast invariance. Our results suggest that a contrast-invariant representation of texture emerges from populations of neurons which are not only selective for naturalistic structure, but are also diverse in their tuning. We conclude that disentangling the representation of multiple variables from a neuronal population requires populations with heterogeneous tuning for the variables of interest.

**Table 1.** Standardized partial regression coefficients (*β*) for each predictor, by area.

| Area | Mean contrast | SD contrast | Mean texture | SD texture |
| --- | --- | --- | --- | --- |
| V1 | -0.38 | 0.08 | 0.13 | **0.47** |
| V2 | -0.04 | -0.20 | 0.02 | **0.67** |
| V4 | 0.01 | 0.18 | 0.03 | **0.49** |

**Fig. 5.**
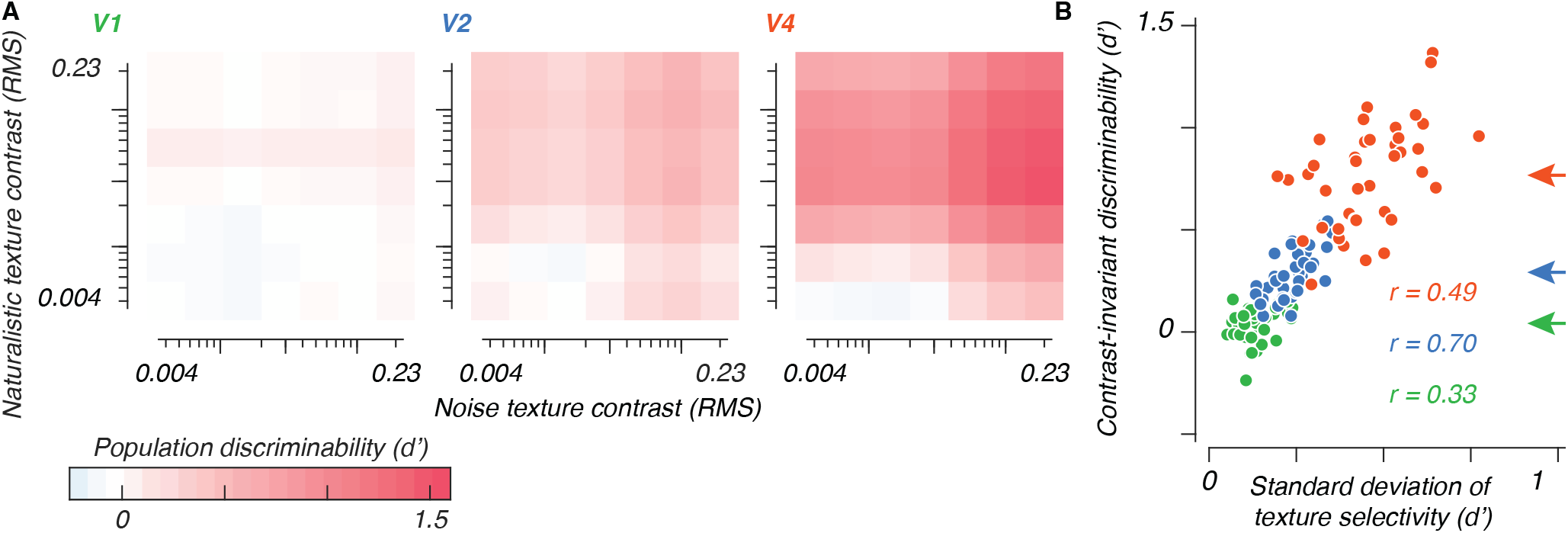
Heterogeneous texture selectivity improves population contrast invariance. A: Mean contrast-independent naturalistic-noise texture discriminability, based on populations of 8 sites (down from 32 in Fig. 4C). As before, the prevalence of red in the V2 and V4 heatmaps is indicative of contrast-invariant naturalistic texture classification. B: Contrast-invariant discriminability for each subsample, versus the standard deviation of texture selectivities for that sample. Each point represents one of 40 population subsamples. Arrows show the contrast-invariant discriminability taken from the average heatmaps in A.

## Discussion

To understand how neural populations might encode separate stimulus dimensions, we measured how the responses of neurons and neural populations are modulated by changes in contrast and naturalistic structure. Contrast modulates activity in V1, V2, and V4 (14–19). Naturalistic structure modulates activity in V2 and V4 (6–10, 20), but not in V1. By varying both dimensions together, we were able to compare contrast sensitivities across areas, and to establish how the encoding of naturalistic structure in V2 and V4 is impacted by changes in contrast. We found that contrast sensitivities were higher in V2 than in V1, and higher still in V4. Contrast sensitivities in V2 and V4 were texture-specific: sensitivities were higher for naturalistic textures than for noise. At the population level, we found that while both V2 and V4 represent naturalistic structure, heterogeneous neural tuning for texture in V4 provided a basis for a robust contrast-invariant readout of naturalistic structure.

### Contrast sensitivity for large texture images

Previous studies of contrast sensitivity in the ventral stream typically used stimuli, often gratings, which were adjusted to the preferred size, orientation, movement, and spatial frequency of the single neuron under study (14–19). Instead of optimized gratings, we used large texture images, each containing a wide variety of orientations and spatial frequencies. We also recorded multiunit activity – we did not sort out single units. As shown in Table 2, we found higher contrast sensitivities than prior studies that used grating or Gabor pattern stimuli.

**Table 2.** Median *c*_50_ values, as reported here, and in previous studies. We converted Michelson contrast values to RMS contrast for comparison with our own results. The value reported by Albrecht and Hamilton (14), marked with an asterisk, included neurons recorded from V1 of both macaques and cats. The study of Sani et al.(15), marked with a dagger, used a rectangular bar. Other studies used gratings or Gabor patterns.

| Citation | Areas recorded | $c_{50}$ (RMS contrast) |
| --- | --- | --- |
| Sclar et al. (17) | V1 | 0.12 |
| Albrecht and Hamilton (14)* | V1 | 0.068 |
| Zheng et al. (18) | V1 | 0.066 |
| **This study – Noise** | **V1** | **0.063** |
| **This study – Naturalistic** | **V1** | **0.065** |
| Zheng et al. (18) | V2 | 0.071 |
| **This study – Noise** | **V2** | **0.054** |
| **This study – Naturalistic** | **V2** | **0.044** |
| Williford and Maunsell (16) | V4 | 0.053 |
| Sani et al. (15)<sup>†</sup> | V4 | 0.018 |
| **This study – Noise** | **V4** | **0.039** |
| **This study – Naturalistic** | **V4** | **0.022** |

Our use of wide-band stimuli and multiunit activity may explain this difference. However, the contrast sensitivity of our sites in V4 was similar to that seen by Sani et al. (15), who used sharp-edged bar stimuli (containing energy at many spatial frequencies). In general, while the relationship in sensitivity across areas was consistent, the elevated sensitivity for naturalistic texture is not fully explained, nor is the role of supersaturation, which was more prevalent in V4, but not necessary for contrast-invariant texture discrimination.

### The importance and basis of contrast invariance

Responses at single sites vary in response to multiple dimensions of stimulus variation. It is therefore impossible to decode the cause of a response change without considering a neural population. For example, while responses in V4 are typically enhanced by naturalistic structure, this enhancement could be overcome for most sites by an opposing change in contrast (Fig. 3C). Satisfyingly, after projecting out the axis corresponding to stimulus contrast, V4 populations could robustly discriminate low contrast naturalistic textures from high contrast noise, using a linear decoder (Fig. 4C). In V4, individual site responses varied widely in texture selectivity (Fig. 3B), making it possible to locate a subspace independent of contrast.

In addition to texture, neurons in V4 are often tuned for the boundary curvature of a given stimulus (21–24). This curvature tuning is robust to changes in position (22), scale (24), and color (25). The tuning of V4 neurons for combinations of shape and texture (21), or combinations of shape and blur (26) is largely separable, suggesting that decoders constructed according to the principles we have employed here could be used to represent these and other stimulus dimensions of interest and relevance to behavior.

Invariance is of course not privileged to V4: populations in inferotemporal cortex (IT) encode the identity of visual objects (27, 28), regardless of changes in position, scale, and context. Nor is invariance limited to vision – measurements from somatosensory cortex have documented separation between touch-related variables (29). In motor cortex, the population representation of movement preparation lies in a dimension orthogonal to the representation of the movement itself (30). Finally, population activity in prefrontal cortex can flexibly encode task-relevant stimulus parameters, and disregard nuisance dimensions (31). The results reported here demonstrate how separable representations may manifest along the visual hierarchy, in a manner that may support the ability of downstream areas to flexibly read out information.

## Materials and Methods

### Visual stimuli

As in previous experiments (8), we generated stimuli using the texture model of Portilla and Simoncelli (5), using the methods detailed in Freeman et al. (6). Briefly, we used photographs of repeating patterns, including both natural and artificial objects, to generate synthetic textures. We used each photograph to generate a texture “family”. We first synthesized “noise” textures, which matched the average spectral content of the original image, as measured with steerable pyramid filters (5). We then synthesized “naturalistic” textures, which matched both the outputs of the oriented filters (as with the noise textures), as well as the correlations between filter types. For each texture family, we ran the synthesis process multiple times, starting with different white noise images. The resultant “samples” were all matched in their spectra and statistics, but differed in local spatial detail.

We used textures from 5 families (Fig. 1). We used 5 different samples of naturalistic and noise textures from each family. These 5 families are the same as those used for the behavioral and neurometric sensitivity measurements reported in Lee et al., 2024 (8). The spatial frequency content of these images is concentrated in a lower band than in the other texture images used in that study (by a factor of roughly 2.7). We adjusted contrast in octave steps down from the root-mean-square contrast of the original image. Our image set totaled 400 images: 5 (families) *×* 5 (samples) *×* 7 (contrasts) *×* 2 (naturalistic or noise) = 350 texture images, plus 50 blank images used to estimate baseline firing rates.

We presented stimuli on a gamma-corrected CRT monitor with a mean luminance of 28 cd m-2, a resolution of 1280 by 960 pixels, and a frame rate of 100 Hz. We seated animals in a custom primate chair 114 cm from the monitor, at which distance it subtended 20 by 15 deg.

### Neurophysiology

#### Experimental procedure

We trained two juvenile *Macaca nemestrina* monkeys, both female, to fixate the central 3 deg of a computer monitor. We used an infrared video-based eye tracking camera (Eyelink 1000) to monitor fixation. The display background was gray, and the center was marked with a red square which was 0.1-0.2 deg across. Texture images appeared after 160 ms in a pseudorandom order. We presented blocks of 4 images, and showed each image for 200 ms, followed by a 200 ms blank interval.

All texture images subtended 6.4 deg and had the same mean luminance as the gray background. Receptive fields were centered within 1.5 deg of the center of gaze, so we centered stimuli at the center of the monitor. The visual stimuli covered the aggregate receptive fields of all recording sites.

We rewarded animals for holding fixation through a block with a juice reward. If an animal broke fixation during a stimulus, we interrupted presentation and presented the interrupted stimulus again later in the overall sequence. We recorded 10 repetitions of each stimulus.

Our stimuli were of course self-similar over space, but given the difference in receptive field sizes between the 3 areas (32), we wondered whether fixational eye movements may have influenced our results. To check, we split our data based on the visual angle the eyes traversed during each stimulus presentation, and recomputed the naturalistic-noise texture selectivity of each site for presentations with below- and above-median traverses (cf. Fig. 3A). We found no evidence that the amount of eye movement on a given stimulus presentation impacted texture selectivity.

We performed all animal procedures in accordance with the National Institutes of Health *Guide for the Care and Use of Laboratory Animals* (33), and with the approval of the New York University Animal Welfare Committee.

#### Neural recording

We recorded multiunit neural responses at the ages of 32 wk (animal M1) and 30 wk (animal M2). In these same animals we have previously shown that the encoding of naturalistic structure is mature by this age (8)). The experiments reported in this paper were conducted as part of the same experimental series as those of Lee et al. and Rodríguez Deliz et al. (8, 32). In each animal, we recorded responses to these stimuli in four separate sessions (three on adjacent days, the fourth was 28 weeks later); the data reported here are from the first session in each animal, but the results were qualitatively similar across sessions. Results based on these sessions are shown in the *SI Appendix* (Figs. S2-S4, which show versions of Figs. 2C, 3C, and 4C, made from data taken during these other sessions). We elected not to combine data from multiple sessions because it is impossible to be certain that each site’s signals represent the same underlying neuronal population from day to day, creating a potential confound for classification analysis. For population analysis, we opted to combine data across animals, to increase the possible range of population sizes. We analyzed data separately by animal, and found qualitatively similar results between the two animals (Figs. S5-S6).

Details of array placement, and receptive field locations can be found in preceding papers (see (32), Fig. 2A, B). In short, we implanted two 96-electrode (“Utah”) recording arrays into the left hemispheres of each animal under sterile conditions. Electrodes were arranged in a 10 *×* 10 square (four positions were not used for recording), had shanks 1 mm long and had an interelectrode spacing of 0.4 mm. Following implantation, we used anatomical landmarks (including gross anatomical landmarks such as sulci and vasculature, which we observed surgically and related to previously documented area boundaries (34, 35)), physiological response properties (e.g. response latencies), and receptive field characteristics to determine the cortical location of array sites. All sites on an array were typically within a single visual area, with one exception, where the array lay on the border between V1 and V2. For the session reported here, we used a total of 58 responsive sites from V1 (from animal M1), 120 sites in V2 (34 from M1 and 86 from M2), and 108 sites in V4 (86 from M1 and 22 from M2).

We recorded bandpass filtered (250 Hz to 7.5 kHz) electrical activity, at a sampling rate of 30 kHz. To minimize the influence of common-mode signals, we subtracted the median sample-by-sample voltage across all sites, and then defined spike events (threshold crossings) as negative voltage deviations at least 3 times the root-mean-squared deviation of the baseline voltage.

### Physiological data analysis

#### Single-site analysis

For analysis, we binned multiunit threshold crossings into 10 ms windows. To determine single-site response latencies, we separately measured the discriminability between blank stimuli, and either naturalistic texture or noise texture stimuli (at full contrast) as a d’ between response distributions: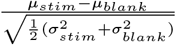, where *µ* reflects the mean of a distribution, and *σ* its standard deviation. We defined the response onset time for a given site as the first deviation above a d’ of *±* 0.4, as long as the following 2 windows were also suprathreshold (and of the same sign). We then computed the median onset time across sites in a given area. For texture analyses, we summed threshold crossings across the 200 ms window starting from these population median response onset times.

To determine whether a site was visually responsive in a given recording session, we combined two metrics. We measured the stimulus-blank discriminability (as a d’) using the summed 200 ms response. We also measured the average response separately for even and odd repetitions of each stimulus. We measured the Pearson correlation between these vectors. We used sites with both a discriminability of 0.1 or greater, and a correlation of 0.4 or greater (the results reported here remained stable across a variety of different metrics and thresholds).

We measured receptive field locations and sizes by recording neural responses to a small spot, which was tiled across the visual field. We used d’ values, summed for 200 ms following response onset, between spot stimuli and blanks to estimate receptive field centers and sizes. All receptive fields were centered roughly 1.5 deg from the center of gaze (below and right of the horizontal and vertical meridian, respectively).

As a single site measure of naturalistic texture discriminability, we measured the d’ between naturalistic and noise texture evoked response distributions.

To model the relationship between stimulus contrast and singlesite response magnitude, we measured the average response to all stimuli at a given contrast, which we fit using the supersaturating model of Peirce (11):

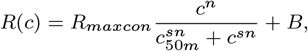

where *R* is the response to a given contrast, *Rmaxcon* is the fit asymptotic value (not necessarily the maximum overall response) of the site. Both *n* and *s* are fit exponents, and *B* is the fit baseline response. This model is an adaptation of the earlier Naka-Rushton model (36). Peirce’s principal innovation is the parameter *s*, which enables the model to fit supersaturating contrast-response data. The presence of the *s* parameter means that the *c*_50*m*_ and *R*_*maxcon*_ parameters do not refer to the half-maximum contrast, nor the maximum response rate, as in the Naka-Rushton model. Here, we refer to them as *c*_50*m*_ and *R*_*maxcon*_ to indicate that the former parameter is only relevant for the model (the m in *c*_50*m*_), while the latter refers to the response at maximum contrast, not necessarily the overall maximum response. For our reported half-maximum contrast values, we parametrically extracted the actual contrast value, referred to as *c*_50_, which represents the contrast eliciting a half-maximum response.

#### Population analysis

To measure population discriminability, we used a training set *M*_*train*_, containing responses to naturalistic and noise texture images from all texture families, to compute linear classifiers. A linear classifier is a hyperplane 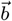 that separates responses by class in *M*_*train*_. To compute these classifiers, we used a linear discriminant, given by the weighted sum of responses from all sites that best separates two response distributions defined by a stimulus feature.

Each column in the *M*_*train*_ matrix contains data from a single recording site, each row corresponds to a response to an individual stimulus presentation, each entry contains the response of a given site on that trial. To measure discriminability we projected held out data *M*_*test*_, onto the orthogonal axis as *M*_*test*_ 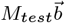. We measured these distances in d’ units. We recorded responses to 5 texture samples for each texture family and contrast. For cross-validation, we used 4 of these samples in *M*_*train*_, and the fifth in *M*_*test*_. We repeated this five-fold cross-validation using all 5 samples.

We measured performance across a number of population sizes, octave-spaced from 2 to 64. Because different recording arrays yielded different numbers of sites, we diagonalized the covariance matrix to remove the impact of correlations between sites. For the results reported in Figure 4, we used a population of size 32, the largest size that allowed us to compare results across areas. Our results were robust across the other population sizes. To estimate variability, for each population size *n*_*pop*_, we used 40 random subsamples of size *n*_*pop*_, drawn from the total sample of visually responsive sites in that area. For the results reported in Figure 5, we used populations of size 8.

We trained two classifiers: one to discriminate naturalistic textures from noise, using responses to all contrasts, and another to discriminate higher contrast textures from lower contrasts, using responses to both naturalistic textures and noise. This yielded two linear axes in the population response space: a naturalistic-noise axis, and a contrast axis. We then used Gram-Schmidt orthogonalization to compute a third axis, a naturalistic-noise axis which is independent of contrast. We measured discriminability between naturalistic textures and noise using both the original naturalistic-noise axis, and the contrast-independent axis (Fig. 4 B and C, Fig. 5). As a metric of “contrast-invariant discriminability,” we took the mean of the naturalistic-noise discriminability values for which the naturalistic texture contrast was lower than the noise texture contrast (i.e. cells below the diagonal of the heatmaps in Figs. 4 and 5).

#### Statistical testing

Where necessary, we used bootstrap resampling to estimate confidence intervals. We performed permutation ANOVAs, after Anderson (12), to measure area-level effects. When significant, we then performed pairwise permutation tests to compare effects either between or within areas.

To estimate the degree to which contrast-invariant discriminability could be attributed to different single-site variables, we performed a multiple regression analysis (13). We computed the standard partial regression coefficients, *β*, using the model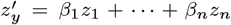, where each *β* represents the increase in contrast-invariant performance, 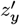 for a change in a given predictor *zn*. Because units are all standard scores, we compare *β* values directly to evaluate how each predictor relates to contrast invariance. For our predictors, we used the mean and standard deviation of contrast and texture selectivity across the 8 sites in a given subsample (four predictors in total).

## ACKNOWLEDGMENTS

We are grateful to members of the Visual Neuroscience Laboratory for advice and discussion. R.T. Raghavan helped with model fitting, Justin Lieber provided useful insight about population analysis. This work was supported by grants from the National Institutes of Health: R01EY022428 (JAM), R01EY024914 (LK, JAM), R01EY031446 (NJM), F31EY031249 (GML), and F31EY031592 (CLRD).

