## Supplemental Figures for "Emergence of a contrast-invariant representation of naturalistic texture in macaque visual cortex"

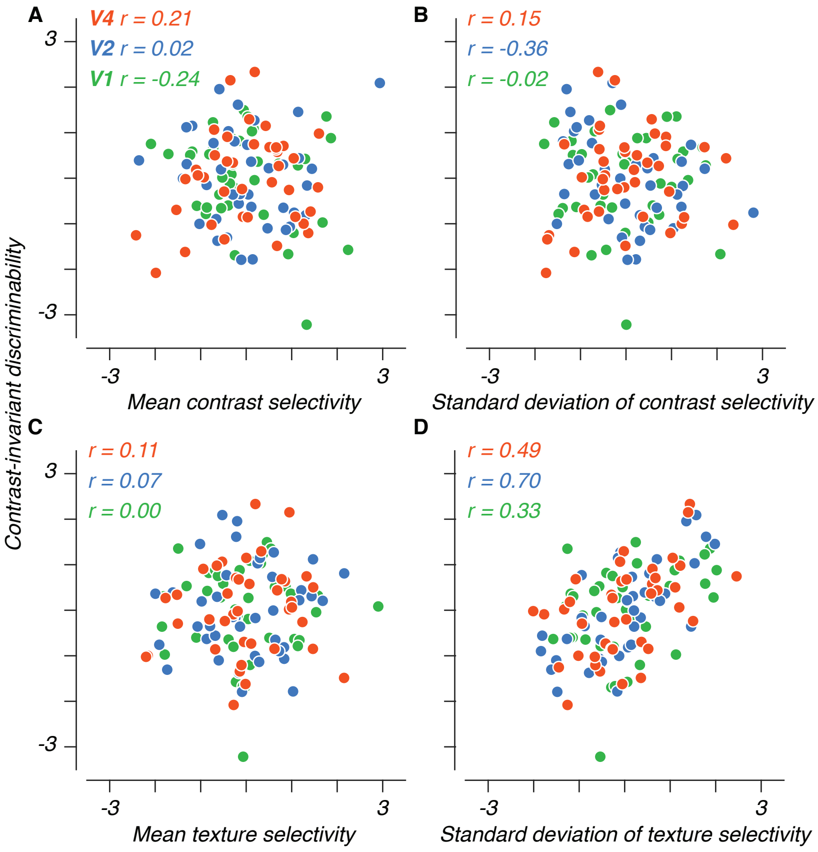


**Fig. S1.** Standardized (Z-scored) plots showing the relationship between contrast invariant discriminability and the first two moments of single site contrast and texture selectivity. Each point shows values taken from one population subsample of 8 sites. A: Performance versus mean contrast selectivity. B: Performance versus the standard deviation of contrast selectivity. C: Performance versus mean texture selectivity. D: Performance versus the standard deviation of texture selectivity (the same data as in Fig. 5B, but Z-scored).

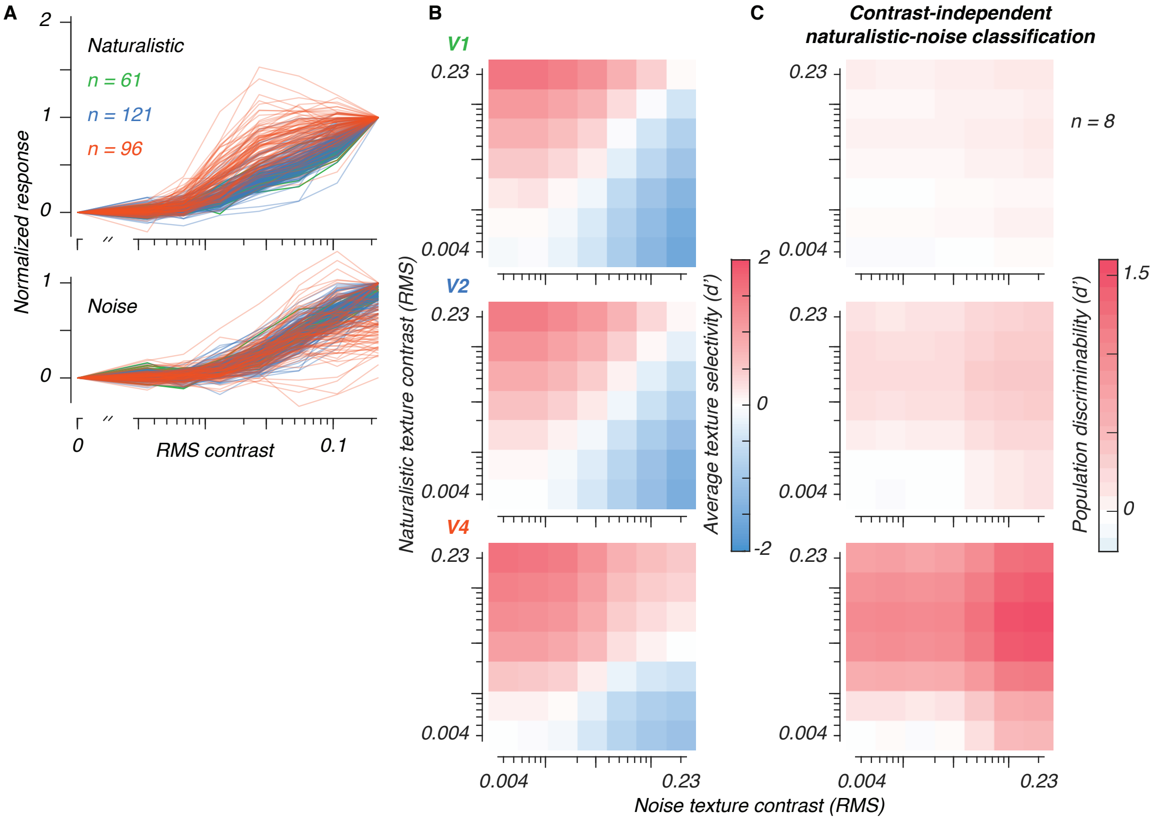


**Fig. S2.** Data taken from the same animals the day after the session used for the figures in the main text. A: Contrast responses to texture stimuli for all sites (as in Fig. 2C). The upper panel shows responses to naturalistic textures, the lower panel shows responses to noise textures. Both sets of curves are normalized to the largest response at maximum contrast for each site, whether it was evoked by naturalistic or noise stimuli. B: Mean single site texture selectivities, measured between responses to naturalistic and noise textures for all possible contrast pairs (as in Fig. 3C). C: Mean naturalistic-noise texture discriminability, based on populations of 32 sites (as in Fig. 4C). Contrast-invariant (below-diagonal) performance was 0.14 in V1, 0.38 in V2, and 1.6 in V4 (for comparison, in Fig. 4C these values were 0.11, 0.45, and 1.6, respectively).

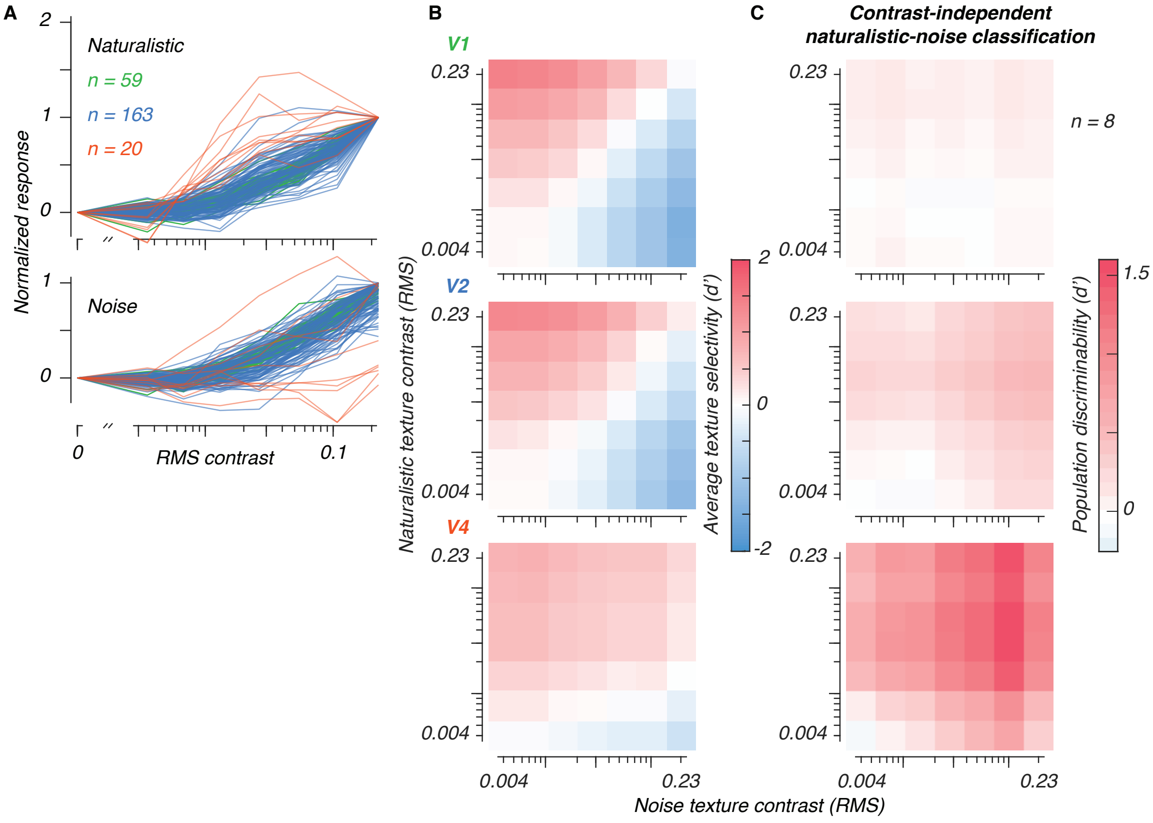


**Fig. S3.** Data taken from the same animals two days after the session used in the main text. Conventions are as in Fig. S1, though note the use of 8-site populations in C (due to the small number of sites recorded in V4). Performance was qualitatively similar to data taken during other sessions. Contrast-invariant (below-diagonal) performance was 0.06 in V1, 0.28 in V2, and 0.86 in V4 (for comparison, in Fig. 5A, which also used populations of 8 sites, these values were 0.04, 0.29, and 0.77, respectively).

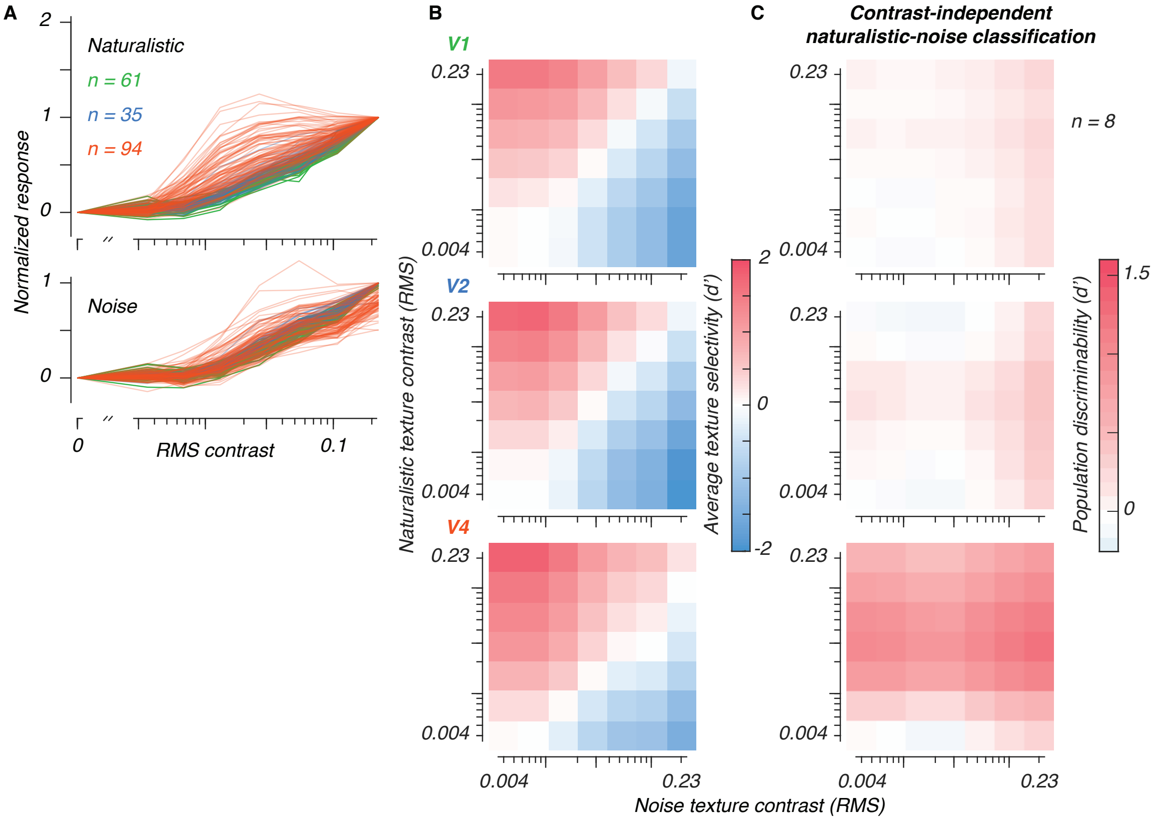


**Fig. S4.** Data taken from the same two animals 28 weeks after the session used in the main text. Conventions are as in Fig. S1, though note the use of 8-site populations in C (due to the low number of sites in V2). Performance in V2 was reduced, as we documented in Supplementary Figure 6 of Lee et al., 2024. The ordinal relationship between areas was nevertheless consistent with data taken during earlier sessions. Contrast-invariant (below-diagonal) performance was 0.12 in V1, 0.18 in V2, and 0.63 in V4 (for comparison, in Fig. 5A, which also used populations of 8 sites, these values were 0.04, 0.29, and 0.77, respectively).

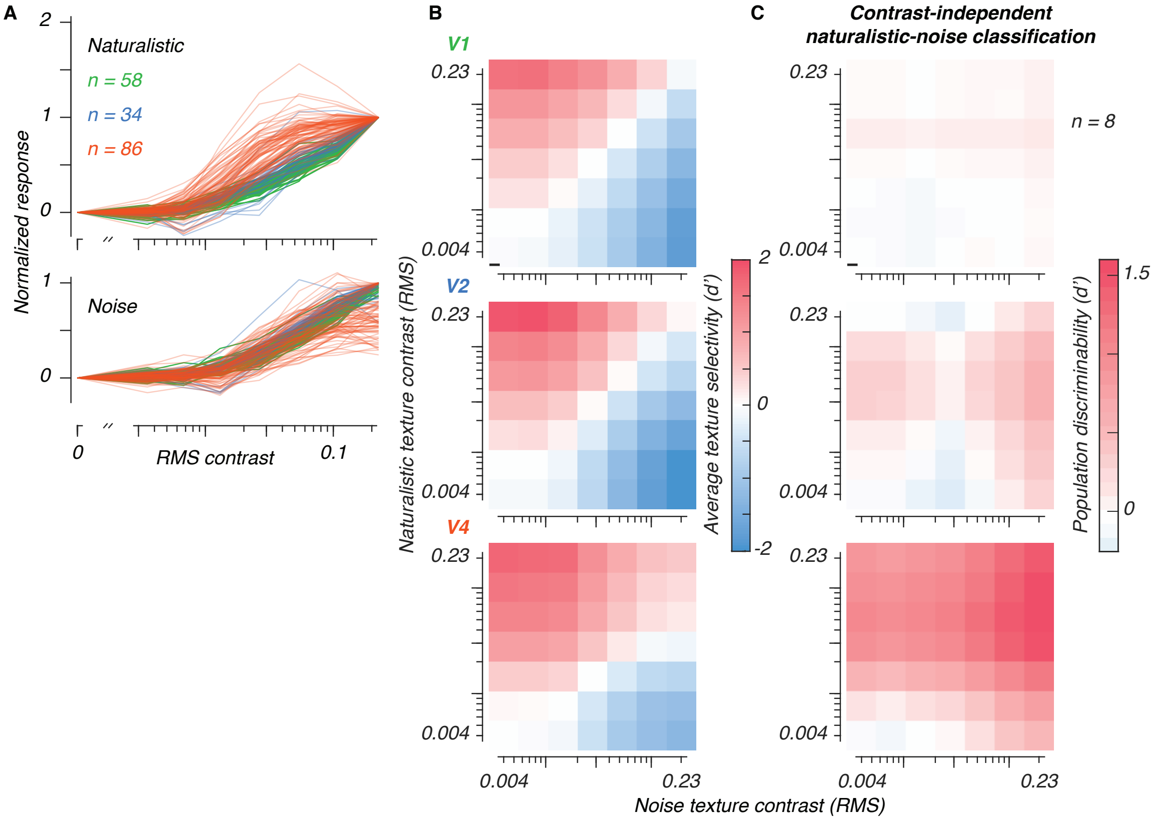


**Fig. S5.** Data taken from animal M1, from the session reported in the main text. Conventions as in Fig. S3, including the use of 8-site populations in panel C. Contrast-invariant (below-diagonal) performance was 0.04 in V1, 0.20 in V2, and 0.78 in V4 (for comparison, in Fig. 5A these values were 0.04, 0.29 and 0.77, respectively).

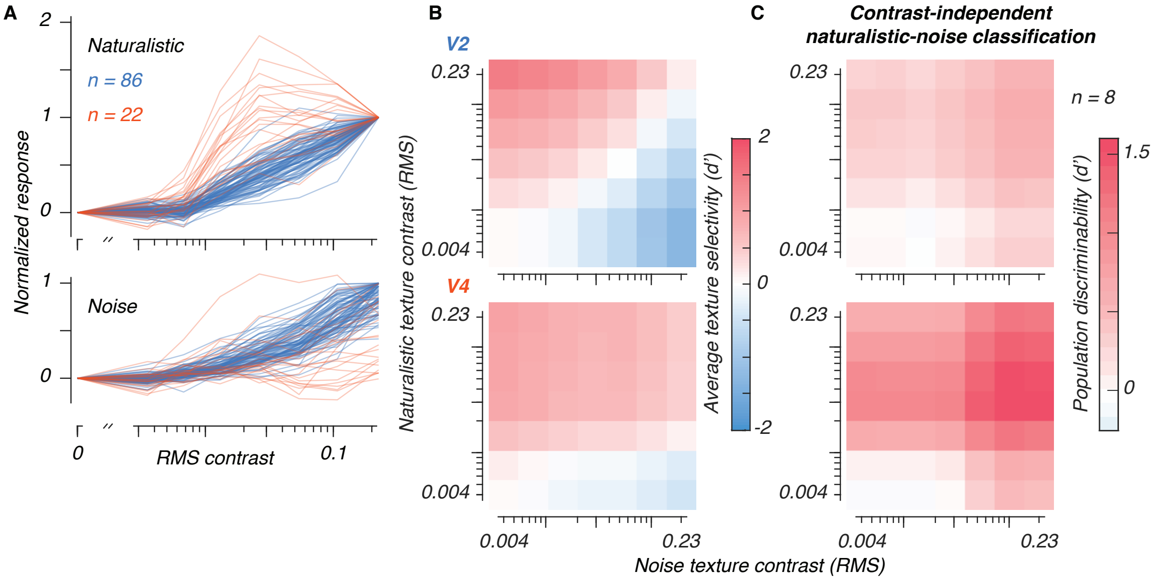


**Fig. S6.** Data taken from animal M2, from the session reported in the main text. Conventions as in Fig. S3, including the use of 8-site populations in panel C. Contrast-invariant (below-diagonal) performance was 0.34 in V2, and 0.81 in V4 (for comparison, in Fig. 5A these values were 0.29 and 0.77, respectively).
